# Electro-Metabolic Sensing Through Capillary ATP-Sensitive K^+^ Channels and Adenosine to Control Cerebral Blood Flow

**DOI:** 10.1101/2021.03.12.435152

**Authors:** Maria Sancho, Nicholas R. Klug, Amreen Mughal, Thomas J. Heppner, David Hill-Eubanks, Mark. T. Nelson

## Abstract

The dense network of capillaries composed of capillary endothelial cells (cECs) and pericytes lies in close proximity to all neurons, ideally positioning it to sense neuro/glial-derived compounds that regulate regional and global cerebral perfusion. The membrane potential (V_M_) of vascular cells serves as the essential output in this scenario, linking brain activity to vascular function. The ATP-sensitive K^+^ channel (K_ATP_) is a key regulator of vascular V_M_ in other beds, but whether brain capillaries possess functional K_ATP_ channels remains unknown. Here, we demonstrate that brain capillary ECs and pericytes express K_ATP_ channels that robustly control V_M_. We further show that the endogenous mediator adenosine acts through A_2A_ receptors and the G_s_/cAMP/PKA pathway to activate capillary K_ATP_ channels. Moreover, K_ATP_ channel stimulation *in vivo* causes vasodilation and increases cerebral blood flow (CBF). These findings establish the presence of K_ATP_ channels in cECs and pericytes and suggest their significant influence on CBF.

**HIGHLIGHTS:**

- Capillary network cellular components—endothelial cells and pericytes—possess functional K_ATP_ channels.
- Activation of K_ATP_ channels causes profound hyperpolarization of capillary cell membranes.
- Capillary K_ATP_ channels are activated by exogenous adenosine via A_2A_ receptors and cAMP-dependent protein kinase.
- K_ATP_ channel activation by adenosine or synthetic openers increases cerebral blood flow.

## INTRODUCTION

Normal brain function requires that every neuron receive an appropriate supply of oxygen, energy metabolites and other nutrients via the cerebral vasculature. Cerebral blood flow (CBF) through the interconnected vascular network of surface arteries, penetrating arterioles, and the vast, complex mesh of capillaries is tuned by the precise and orchestrated action of neurovascular coupling mechanisms, which link blood flow to the metabolic needs of neurons. Capillaries account for a large majority of this vascular network and critically cover all brain regions (Gould et al., 2017), ideally positioning the capillary endothelial cells (cECs) and pericytes that constitute capillaries—and the astrocytic endfeet that completely cover them—in close contact with all neurons in the brain. In fact, we recently demonstrated that cECs are exquisitely capable of sensing extracellular K^+^, a byproduct of neural activity, via inward-rectifier K^+^ (Kir2.1) channels and responding by initiating a retrograde hyperpolarizing (electrical) signal that propagates upstream, causing dilation of feeding arterioles and thereby directing blood flow to the active brain region that initiated the signal (Longden et al., 2017).

Adenosine, a signaling nucleotide that arises via hydrolysis of astrocyte-released ATP (Dunwiddie et al., 1997; Zhang et al., 2003) or direct release from neurons/glial cells upon certain pathological conditions (e.g., hypoxia or ischemia) (Park et al., 1992; Latini et al., 1999; O’Regan, 2005), is a potent endogenous vasodilator that plays a central role in matching blood distribution to tissue metabolic demands (Berne, 1980). In many vascular systems, including the mesenteric circulation, the vasodilatory actions of adenosine are partly attributable to activation of ATP-sensitive potassium (K_ATP_) channels via G_s_-protein–coupled A_2A_ receptors (A_2A_R). The resulting stimulation of adenylyl cyclase (AC) raises intracellular levels of cyclic AMP (cAMP), which binds and activates cAMP-dependent protein kinase (PKA) to ultimately phosphorylate multiple sites in the K_ATP_ channel complex and promote its opening (Kleppisch and Nelson, 1995; Quayle et al., 1997; Li and Puro, 2001; Quinn et al., 2004). The resulting K_ATP_ channel-mediated K^+^ efflux hyperpolarizes the cell membrane, closing voltage-gated Ca^2+^ channels and thereby inducing vessel dilation (Standen et al., 1989; Nelson et al., 1990).

K_ATP_ channels are potent regulators of the membrane potential (V_M_) of vascular cells. In smooth muscle cells (SMCs), K_ATP_ channels are octamers of four pore-forming Kir6.1 subunits and four accessory sulfonylurea receptors (SUR2B) (Li et al., 2013). Vascular K_ATP_ channel function is highly sensitive to the relative intracellular concentrations of ATP and ADP ([ATP]/[ADP] ratio) (Quayle et al., 1994). These channels are also stimulated by synthetic K^+^ channel openers (e.g., pinacidil) or endogenous vasodilators, including adenosine, vasoactive intestinal peptide and calcitonin gene-related peptide (CGRP), and are inhibited by oral hypoglycemic sulfonylurea drugs such as glibenclamide (GLIB) (Standen et al., 1989; Nelson et al., 1990; Nelson and Quayle., 1995; Quayle et al., 1997; Zhang et al., 1994), an antagonist of K_ATP_ channels whose selectivity has been well established in a variety of cell types, including pancreatic β-cells, cardiac myocytes and vascular SMCs (Sturgess et al., 1985). In the brain, however, SMCs may not be ideal sensors of endogenous vasoactive agents, specifically adenosine, due to the relatively low density of SMC-containing arterioles within the cortex. In contrast, neuron-juxtaposed brain cECs and pericytes, especially the latter, robustly express the genes encoding A_2A_R (*Adora2a*) and the vascular K_ATP_ channel subunits Kir6.1 (*Kcnj8*) and SUR2B (*Abcc9*) (Vanlandewijck et al., 2018; Sabbagh et al., 2018; Zhang et al., 2014), suggesting that adenosine could act through capillary cell K_ATP_ channels to control CBF.

Here, we demonstrate that the main cellular components of brain capillaries—ECs and pericytes—express functional K_ATP_ channels that strongly influence resting V_M_. We further show that capillary K_ATP_ channels respond to extracellular adenosine, a key neuron/glial signaling molecule, via the A_2A_R-G_s_-PKA pathway. Moreover, activation of K_ATP_ channels by adenosine or pinacidil *in vivo* increases CBF. Collectively these findings establish the presence of functional K_ATP_ channels in the brain capillary network and demonstrate their potential physiological impact on brain hemodynamics.

## RESULTS

### Brain Capillary Cells Express Functional K_ATP_ Channels

To ascertain whether brain capillary networks express functional K_ATP_ channels, we employed patch-clamp electrophysiology on freshly isolated cEC and pericytes. cECs, which are easily recognized by their characteristic tube-like shape, exhibited pronounced linear inward Kir2.1 currents at hyperpolarized potentials (K^+^ equilibrium potential [E_K_] = −23 mV at 60 mM external K^+^) (**Figure 1A**, *black inset*). The identification of native pericytes was facilitated using NG2-DsRed-BAC transgenic mice, which express the DsRed fluorescent protein under control of the NG2 promoter encoding the putative pericyte marker, neural/glial antigen 2 (NG2). Freshly isolated pericytes exhibited characteristic protruding cell bodies and thin-stranded projections. We found that the current-voltage relation of pericytes is unique and distinct from that of ECs and SMCs. When bathed in 60 mM K^+^, whole-cell currents in pericytes exhibited a non-linear inward current and a modest outward current that was less than 1/3 that of the SMC outward current at +40 mV (**Figure 1A**, *red inset*; **S1. Related to Figure 1**). In contrast, cECs showed minimal outward currents positive to E_*K*_.

**Figure 1.**
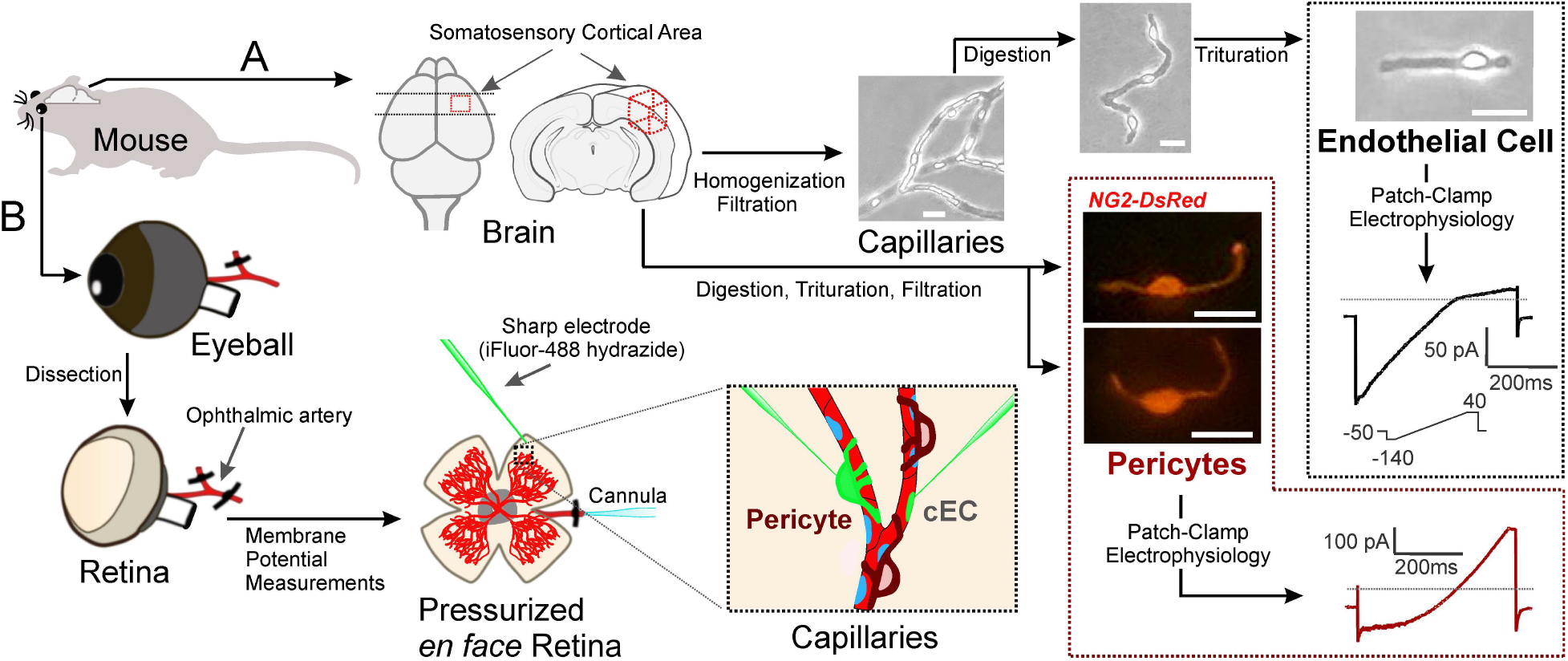
Experimental paradigm used to explore the expression of functional K_ATP_ channels in the capillary network. (A) Schematic depiction of brain cEC and pericyte isolation for patch-clamp analysis. *Left to right*: A small piece of brain somatosensory cortex is mechanically disrupted and filtered to yield capillary segments. Single capillary cells are released after enzymatic digestion and trituration. Pericyte identification is aided by the use of NG2-DsRed-BAC transgenic mice. Endothelial cells and pericytes exhibit different electrophysiological profiles (*black and red trace*, respectively). (B) Ilustration of the intact pressurized retina preparation and measurement of membrane potential *in situ* using sharp microelectrodes. The ophthalmic artery of an isolated mouse retina is cannulated, and the retina tissue is pinned down *en face*, allowing visualization of the entire superficial microvasculature and facilitating the impalement of a cell of interest (cEC or pericyte; *inset*). The phenotype of the impaled cell is confirmed by including iFluor-488-conjugated hydrazide in the glass sharp electrode. Scale bars, 10 μm.

Using the conventional whole-cell configuration of the patch-clamp technique, we tested whether the potent K_ATP_ channel opener pinacidil evokes K_ATP_ currents in isolated brain cECs and pericytes. Cells were held at a membrane potential of −70 mV, bathed in a 60 mM K^+^ solution and dialyzed with a 140 mM K^+^ intracellular solution. Under these experimental conditions, K^+^ currents are inward (Quayle et al., 1994). In cECs dialyzed with a low ATP concentration (0.1 mM) and a relative high ADP concentration (0.1 mM), pinacidil (10 μM) increased whole-cell currents by an average of −26.1 ± 2.2 pA/pF (**Figure 2A and 2E**), an effect that was inhibited by subsequent application of glibenclamide (GLIB; 10 μM). PNU-37883A, a putative selective antagonist of vascular K_ATP_ channels, also reversed pinacidil-induced currents in cECs dialyzed with 0.1 mM ATP, reducing them by ~75% (from −23.7 ± 1.1 to −6.1 ± 1.4 pA/pF) (**S2. Related to Figure 2**). Interestingly, pinacidil-evoked, GLIB-sensitive currents in pericytes (−68.4 ± 10.3 pA/pF) were ~2.6-fold greater than those in cECs under identical experimental conditions (**Figure 2B and 2E**). Notably, GLIB-sensitive currents induced by pinacidil were very small in single cerebral pial artery SMCs (−1.6 ± 0.2 pA/pF) and parenchymal arteriole SMCs (−1.2 ± 0.1 pA/pF) dialyzed with 0.1 mM ATP (S1. Related to **Figure 2**). Note that whole-cell currents recorded in a subset of pial SMCs (n = 3) remained unaltered in the presence of pinacidil. These findings suggest that the hyperpolarizing GLIB-sensitive K_ATP_ current induced by pinacidil is initiated almost exclusively in the capillary network.

**Figure 2.**
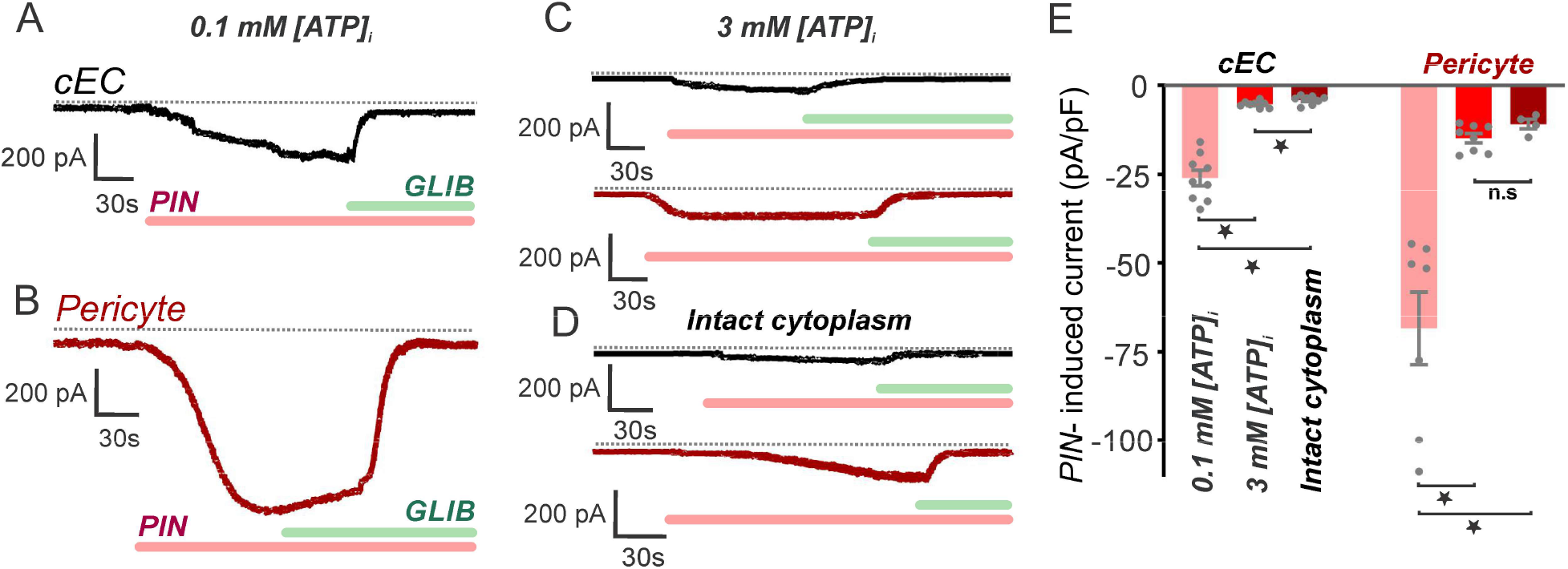
Brain cECs and pericytes express functional K_ATP_ channels. Representative traces illustrating glibenclamide (GLIB; 10 μM)-sensitive currents induced by pinacidil (PIN; 10 μM) from a holding potential of −70 mV in freshly isolated capillary ECs or pericytes. External and internal K^+^ were 60 and 140 mM, respectively. Dotted line represents 0 current level. (A, B) Currents recorded in the whole-cell configuration in capillary ECs (*n* = 9) (A) or pericytes (*n* = 7) (B) dialyzed with 0.1 mM ATP. (C) Currents recorded in the whole-cell configuration in capillary ECs (*top*, black trace) or pericytes (*bottom*, red trace) (*n* = 8 each) dialyzed with 3 mM ATP. (D) Currents recorded from cECs (*top, black trace*) (*n* = 8) and pericytes (*bottom, red trace*) (*n* = 4) in the perforated patch-clamp configuration (cytoplasm intact). For A–C, the pipette solution also contained 0.1 mM ADP in addition to the indicated concentration of ATP. (E) Summary data showing pinacidil-induced current density prior to and following GLIB treatment in cECs and pericytes dialyzed with different ATP concentrations or recorded in the perforated patch configuration. Values are presented as means ± SEM (\**P* < 0.05, paired Student’s *t*-test).

K_ATP_ channel activity is also influenced by the cytosolic [ATP]/[ADP] ratio. Specifically, K_ATP_ channels are activated by a low concentration of cytosolic ATP or by an elevated ADP concentration; thus, an increase in the [ATP]/[ADP] ratio would be expected to inhibit K_ATP_ channel activity. Consistent with this, K_ATP_ currents were ~5-fold smaller (−5.3 ± 0.3 pA/pF; *n* = 8) in cECs dialyzed with 3 mM ATP and 0.1 mM ADP ([ATP]/[ADP] ratio = 30) than those in cECs dialyzed with a 1:1 ratio (0.1 mM each) of ATP to ADP (−26.1 ± 2.2 pA/pF). Similarly, raising the internal [ATP]/[ADP] ratio from 1 to 30 reduced the average K_ATP_ current in pericytes ~4.6-fold (from −68.4 ± 10.3 pA/pF to −14.86 ± 1.33 pA/pF) (**Figure 2C and 2E**). Finally, an examination of whole-cell currents using the perforated-patch configuration, in which the cytoplasmic composition remains undisturbed, revealed that currents evoked by pinacidil in intact cECs and pericytes were smaller (~25% in both cell phenotypes) than those observed following dialysis with 3 mM ATP and 0.1 mM ADP (**Figure 2D and 2E**). These findings collectively support the idea that capillary cells express functional K_ATP_ channels that are activated by pinacidil and inhibited by GLIB, PNU-37883 and ATP. Furthermore, K_ATP_ channel currents in the absence of pinacidil were too small to be detected in the perforated-patch configuration (i.e., with physiological intracellular [ATP]/[ADP] ratios).

### Activation of K_ATP_ causes a pronounced membrane potential hyperpolarization

The role of capillary K_ATP_ channels in controlling membrane potential was assessed *in situ* using our recently developed intact pressurized retina preparation and sharp microelectrodes. In this preparation, the ophthalmic artery of an isolated retina is cannulated and the retina tissue is pinned down en face, allowing visualization of the entire superficial microvascular network and enabling the impalement of cells of interest (**Figures 1B and 3A**). To determine whether we were recording from an cEC or pericyte, we included fluorescent iFluor-488-conjugated hydrazide in the glass microelectrode pipette. The resulting dye-filled cells (green) were identifiable based on their morphology and were subsequently verified in a subset of experiments using NG2-DsRed mouse retinas, in which green dye can readily be visualized filling red pericytes (yellow in merged images) or underlying ECs (**S3. Related to Figure 3**). Note that these maneuvers revealed extensive coupling of cECs to each other, as evidence by dye movement between adjacent cells, but showed little or no coupling between pericytes or between cECs and pericytes (**Figure 3B and 3C**). After cannulating and pressuring (50 mm Hg) the ophthalmic artery, cells were impaled, pinacidil was applied and membrane potential was measured in cECs and pericytes using microelectrodes. In cECs, pinacidil (10 μM) induced a robust membrane hyperpolarization to −74.0 ± 2.4 mV from the steady state V_M_ of −33.6 ± 1.0 mV. Subsequent superfusion with GLIB (20 μM) depolarized endothelial V_M_ to −24.7 ± 1.7 mV (**Figures 3D and 3F**). Similar results were observed with pericytes, where, on average, pinacidil hyperpolarized cells from −40.5 ± 4.4 mV to −78.2 ± 5.8 mV, an effect that was reversed by GLIB (−44.7 ± 0.6 mV) (**Figures 3E and 3F**). These data highlight the presence of K_ATP_ channels in capillary cells and suggest that they play a major role in regulating membrane potential in these cells.

**Figure 3.**
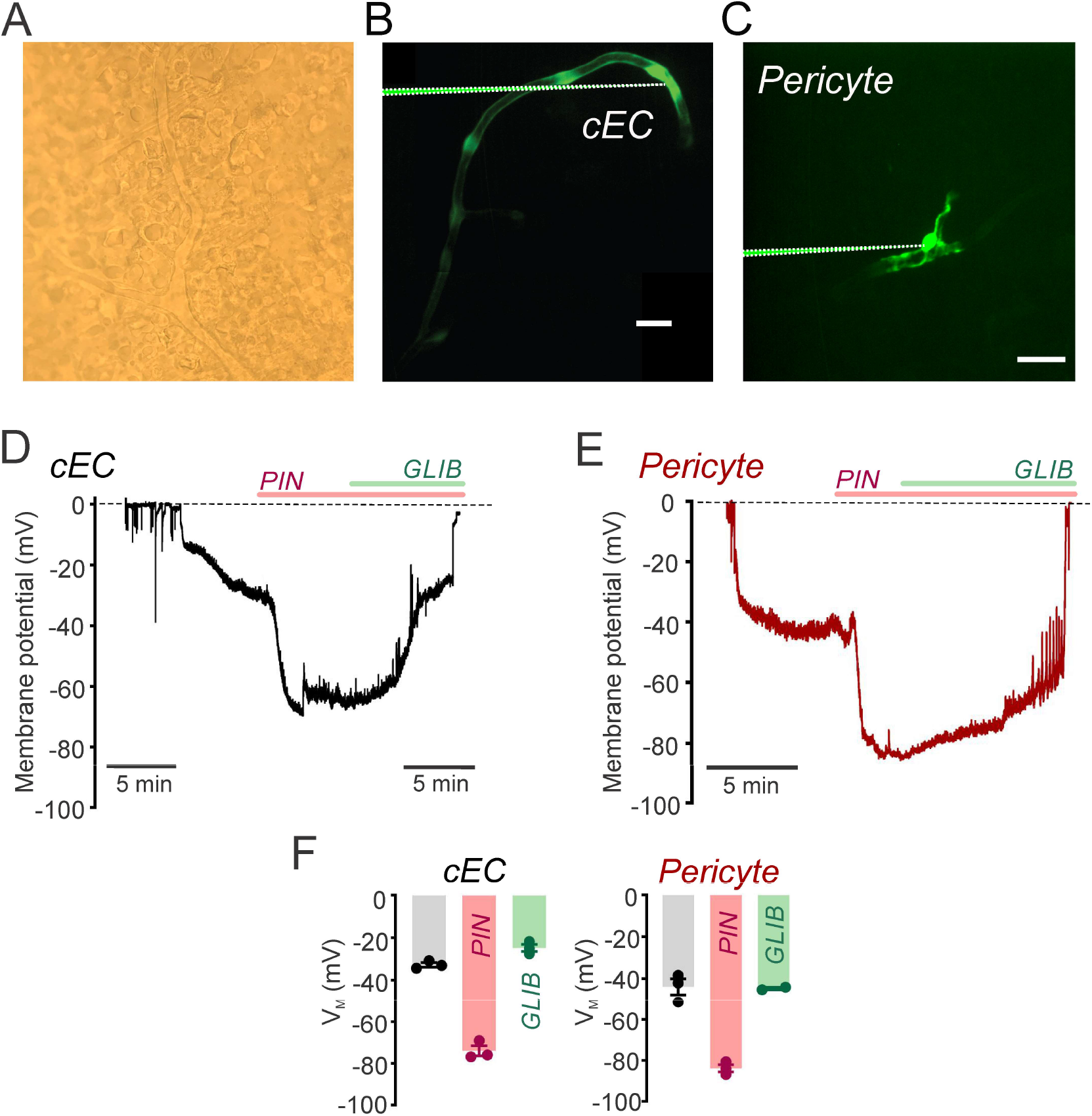
Activation of K_ATP_ channels causes profound membrane potential hyperpolarization. (A) Representative high-magnification, brightfield image of the intact *en face* pressurized retina preparation used for membrane potential measurements. (B) Stitched z-projection of spinning-disk confocal images illustrating an impaled capillary endothelial cell and the diffusion of iFluor-488-conjugated hydrazide to coupled adjacent cells, staining mainly cEC bodies. (C) In pericytes, fluorescent hydrazide labeling remained exclusively in the cell body and processes of the impaled cell. Note that z-projections were generated after removal of the microelectrode; the previously imaged position of the microelectrode was added to the images for better interpretation of impalement location. Scale bars, 20 μm. (D, E) Representative membrane potential recordings from a cEC (*black trace*; n = 3) and a pericyte (*red trace*; n = 3) illustrating the robust hyperpolarization induced by pinacidil (*PIN*; 10 μM) and subsequent block by glibenclamide (*GLIB*; 20 μM). Dashed line represents 0 mV baseline before impalement of the cell. (F) Summary data showing the effects of pinacidil and GLIB on the V_M_ of cECs and pericytes. Data are expressed as means ± SEM (\**P* < 0.05, paired Student’s *t*-test).

### Adenosine Activates Capillary K_ATP_ Channels through A_2A_Rs and PKA

The potent endogenous vasodilator adenosine has been reported to activate K_ATP_ channels via A_2A_Rs and PKA in a variety of vascular systems, including the mesenteric and coronary vasculature (Kleppisch & Nelson, 1995; Dart & Standen 1993) (**Figure 4A**). To test whether brain capillary K_ATP_ channels respond to extracellular adenosine, we recorded currents in native cECs and pericytes dialyzed with a pipette solution containing 0.1 mM ATP, 0.1 mM ADP and 140 mM K^+^ (V_M_ = −70 mV, 60 mM external K^+^). As illustrated in **Figure 4B**, adenosine (25 μM) robustly stimulated whole-cell currents in cECs and pericytes, increasing these currents by an average of −17.2 ± 1.4 pA/pF and −59.5 ± 12.4 pA/pF, respectively. As expected, adenosine-induced currents were significantly larger (~3.5-fold) in pericytes than cECs under the same experimental conditions. In both cases, these currents were inhibited by GLIB (10 μM) and PNU-37883 (10 μM) (S2. Related to **Figure 4**), suggesting that adenosine effects in capillary cells are specifically attributable to activation of K_ATP_ channels. In contrast, whole-cell currents in SMCs (dialyzed with 0.1 mM ATP and 0.1 mM ADP; ATP/ADP ratio = 1) isolated from pial arteries or parenchymal arterioles were unaffected by adenosine or GLIB, whereas inward currents recorded at a holding potential of −70 mV were blocked by subsequent addition of Ba^2+^ at a concentration (100 μM) selective for Kir2 channels (S1. Related to **Figure 4**). GLIB-sensitive currents were also robustly activated in both capillary cell types by the endogenous vasodilator CGRP, which similarly signals through G_s_-coupled GPCRs and cAMP/PKA to activate K_ATP_ channels in the systemic vasculature (Nelson et al., 1990) (**S4. Related to Figure 4**).

**Figure 4.**
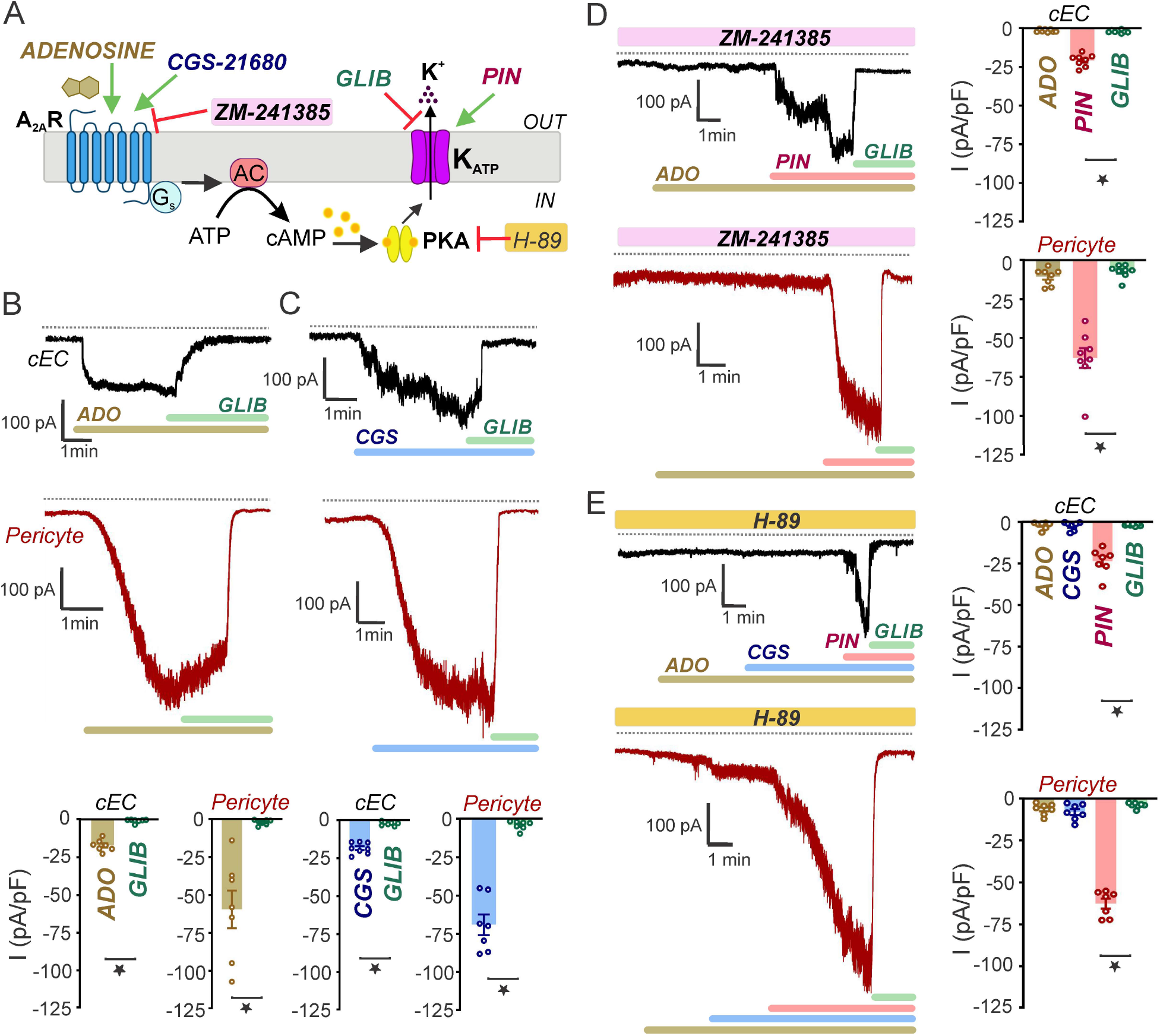
Adenosine activates capillary K_ATP_ channels via A_2A_Rs and PKA. (A) Schematic diagram illustrating the involvement of A_2A_R signaling through the G_s_-AC-PKA intracellular pathway in the activation of K_ATP_ channels. The A_2A_R and PKA agonists/antagonists used are indicated. (B) Representative recordings (*top*) and summary data (*bottom*) showing the stimulatory effects of adenosine (*ADO*; 25 μM) on GLIB (10 μM)-sensitive currents from native cECs (*n* = 7) and pericytes (*n* = 7). (C) GLIB-sensitive currents evoked by the adenosine A_2_ receptor agonist CGS-21680 (*CGS*; 500 nM) in cECs (*n* = 8) and pericytes (*n* = 7). (D) Original traces (*left*) and summary data (*right*) showing effects of adenosine (25 μM) and pinacidil (*PIN*; 10 μM) in cECs (*top*; n = 8) and pericytes (*bottom*; n = 8) treated with the adenosine A_2_ receptor blocker ZM-241385 (30 nM). (E) Inhibition of adenosine- and CSG-21680-induced K_ATP_ currents in cECs (*n* = 7) and pericytes (*n* = 7) by the specific PKA inhibitor H-89 (1 μM) and efficient activation of GLIB-sensitive currents in both cell types by pinacidil. External K^+^ was 60 mM and cells were dialyzed with 0.1 mM ATP, 0.1 mM ADP and 140 mM K^+^. Membrane current was recorded at a V_M_ of −70 mV. Dotted line represents 0 current level. Data are expressed as means ± SEM (\**P* < 0.05, paired Student’s *t*-test).

To explore the mechanisms underlying capillary K_ATP_ channel activation by adenosine, we next tested specific activators and inhibitors of A_2A_Rs, first examining the effect of the selective A_2A_R agonist, 2-4-(2-carboxethyl)-phenethylamino)-5’-N-ethylcarboxamidoadenosine hydrochloride (CGS-21680), on whole-cell K^+^ currents. We found that CGS-21680 (500 nM) induced GLIB-sensitive currents in cECs and pericytes with magnitudes of −18.4 ± 1.3 pA/pF and −69.1 ± 6.7 pA/pF, respectively, values comparable to those measured in response to adenosine exposure (**Figure 4C**). To further analyze the specific contributions of A_2A_R activation to K_ATP_ channel activity, we tested the effect of the potent A_2A_R antagonist, ZM-214385. In capillary ECs or pericytes pretreated with extracellularly applied ZM-214385 (30 nM), adenosine failed to evoke currents, whereas GLIB-sensitive currents induced by pinacidil, which directly opens K_ATP_ channels, were not significantly affected by pretreatment with ZM-214385 in cECs (−20.7 ± 1.4 pA/pF) or pericytes (−62.8 ± 6.3 pA/pF) (**Figure 4D**). Taken together, these findings indicate that adenosine acts via A_2A_Rs to stimulate K_ATP_ channels in brain capillary ECs and pericytes.

In many vascular beds, K_ATP_ activation through A_2A_Rs is mediated by downstream signaling pathways involving stimulation of AC and elevation of intracellular levels of cAMP, which binds and activates PKA (Kleppisch & Nelson, 1995) (**Figure 4A**). To explore coupling of A_2A_Rs in brain capillaries to PKA signaling, we tested the effects of the membrane-permeant PKA inhibitor H-89. Pretreatment of cECs and pericytes with H-89 (1 μM) abolished both adenosine- and CGS-21680–induced GLIB-sensitive currents. Unlike the case for A_2A_R agonists, GLIB-sensitive currents evoked by the K_ATP_ activator pinacidil remained unaltered after H-89 treatment in both cECs (−23.6 ± 3.0 pA/pF) and pericytes (−62.6 ± 2.9 pA/pF) (**Figure 4E**). These data support a role for PKA in activation of K_ATP_ channels in the brain capillary network.

### K_ATP_ Channels Modulate CBF

Finally, to examine the impact of K_ATP_ channel activity in the regulation of CBF, we monitored *in vivo* changes in CBF in the mouse somatosensory cortex using laser Doppler flowmetry in response to surface application of the K_ATP_ agonists pinacidil or adenosine in the absence or presence of GLIB (**Figure 5A**). CBF, measured 30 minutes after superfusion of pinacidil (10 μM), increased by 60.0% ± 3.5%; this effect was significantly blunted by pretreatment with GLIB (10 μM), which reduced the relative increase to 16.6% ± 4.5% (**Figure 5B**). Adenosine (5 μM) superfusion similarly induced an increase in CBF (26.8% ± 2.2%) that was also greatly attenuated by GLIB (relative increase, 7.2% ± 1.1%) (**Figure 5C**). Collectively, these findings demonstrate that activation of capillary K_ATP_ channels increases CBF.

**Figure 5.**
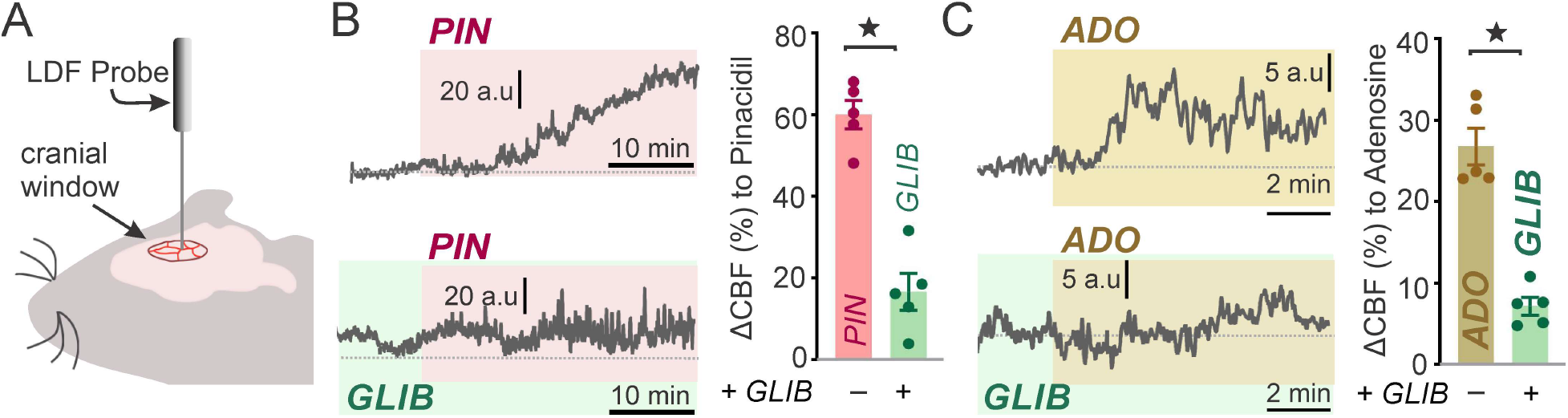
Activation of K_ATP_ channels increases CBF. (A) Experimental setup. CBF responses were continuously monitored in the somatosensory cortex through a closed cranial window using laser-Doppler flowmetry (LDF). (B) Response to pinacidil. Original traces (*left*) and summary data (*right*) illustrating changes in CBF induced by pinacidil (*PIN*; 10 μM) superfusion in the absence or presence of glibenclamide (*GLIB*; 10 μM; *n* = 5 mice/group). (C) Response to adenosine. Representative traces (*left*) and summary data (*right*) showing changes in CBF induced by superfusion of adenosine (*ADO*; 5 μM) in the absence and presence of GLIB (*n* = 5 mice/group). Data are expressed as means ± SEM (\**P* < 0.05, paired Student’s *t*-test).

## DISCUSSION

The dense, anastomosing capillary network represents the primary site of oxygen and nutrient exchange between the vasculature and neurons of the brain. Every neuron in the brain lies within ~10–20 μm of a neighboring capillary, an architectural relationship that favors intimate communication between neurons and capillaries (Attwell et al., 2010). We have previously demonstrated the physiological relevance of this concept, revealing that capillaries act as a neural activity sensory web that responds to K^+^ released during neuronal activity through cEC Kir2.1 channel activation, which causes a retrograde, propagating hyperpolarizing signal that dilates upstream arterioles and increases blood flow to the active neuronal region (Longden et al, 2017). Notably, capillaries are also enwrapped by astrocytes, and are thus similarly poised to respond to astrocyte-derived vasoactive modulators, in particular adenosine, which is constantly released from astrocytes (Haydon & Carmignoto, 2006). One potential end-target of adenosine is the K_ATP_ channel, which has been shown to functionally link A_2A_R-GsPCR signaling to vasodilation via cAMP/PKA in other vascular beds (Quayle et al., 1997). However, whether the K_ATP_ channel is expressed in cells that constitute capillaries is unknown. Recent work from our laboratory has helped to define the biophysical signatures of ion channels in cECs (Longden et al., 2017; Harraz et al., 2018) and elucidate their influence on capillary function; however, the functional ion channel repertoire in these cells still remains incompletely understood. These gaps in our knowledge are even more striking in the case of native brain pericytes, where ion channel expression and function have remained largely uninvestigated. In the present study, we explored the presence of K_ATP_ channels in capillary cells and assessed their potential impact on brain vascular function. Our electrophysiological findings demonstrated that brain capillary cells—both cECs and pericytes—possess K_ATP_ channels that operate as key determinants of membrane potential. Direct electrophysiological recordings from single pericytes freshly isolated from the brain capillary network revealed current-voltage relationships distinct from those of other vascular cells, characterized by a robust, nonlinear, inward K^+^ current at hyperpolarized potentials and an outward current at depolarized potentials that is less prominent compared with that in SMCs (**Figure 1**). We further found that the endogenous vasoactive agent adenosine activates K_ATP_ channels in the capillary network by activating the A_2A_R-G_s_-AC-PKA signaling cascade. Lastly, an *in vivo* analysis of CBF demonstrated robust vasodilatory effects following capillary K_ATP_ channel stimulation. Collectively, our data provide multiple lines of evidence supporting the functional importance of the K_ATP_ channel in brain capillaries and underscore its potential to serve as a physiological bridge between neuronal/astrocyte activity and vascular function and thereby profoundly impact cerebral hemodynamics.

ATP-sensitive K^+^ channels play a central role in setting the membrane potential of vascular smooth muscle cells (Standen et al., 1989; Quayle et al., 1997). K_ATP_ channel opening increases K^+^ efflux, resulting in hyperpolarization, closure of voltage-gated Ca^2+^ channels, and subsequent vessel dilation (Standen et al., 1989; Nelson et al., 1990, Knot and Nelson, 1995). K_ATP_ channels are activated by a variety of K^+^ channel openers, including pinacidil, cromakalim and diazoxide, among others. These channels are also inhibited by oral hypoglycemic drugs, such as glibenclamide and tolbutamide (Quayle et al., 1995; Quayle et al., 1997). Notably, existing single-cell RNA-sequencing data for mouse brain vascular cells indicate that both cECs and pericytes—especially the latter— express the genes *Kcnj8* and *Abcc9* encoding the K_ATP_ channel vascular subunits Kir6.1 and SUR2B, respectively (Vanlandewijck et al., 2018; Sabbagh et al., 2018; Zhang et al., 2014). Pinacidil evoked robust GLIB-sensitive K^+^ currents in both cECs and capillary pericytes dialyzed with a low (1:1) [ATP]/[ADP] ratio (0.1 mM ATP and 0.1 mM ADP) concentrations, but pinacidil-induced currents were markedly reduced in cells dialyzed with a higher (30:1) [ATP]/[ADP] ratio or recorded under conditions (perforated patch-clamp configuration) that maintain cytoplasmic components intact (**Figure 2**). Our findings are in line with previous electrophysiological work performed in capillaries isolated from the heart (Mederos y Schnitzler et al., 2000) and in retinal pericytes (Li and Puro, 2001; Wu et al., 2006). Pinacidil-evoked K^+^ currents were also blocked by PNU-37883, a well-established selective inhibitor of vascular K_ATP_ channels (Wellman et al., 1999). Collectively, these data provide convincingly support for the idea that brain cECs and pericytes possess functional K_ATP_ channels with pharmacological properties similar to those of other smooth muscle K_ATP_ channels (Quayle et al 1997).

Our recently developed intact pressurized retina preparation (Gonzales et al., 2020; Ratelade et al., 2020) provides the means for confirming the presence K_ATP_ channels and experimentally test their contribution to membrane potential in intact capillaries under physiological pressure and flow conditions and within a native environment that reflects their central nervous system context. Using sharp microelectrodes, we demonstrated that activation of K_ATP_ channels with pinacidil robustly hyperpolarized the membrane potential of both cECs and pericytes, effects that were reversed by application of GLIB (**Figure 3**). These findings are comparable to previous results obtained from impaled SMCs in pressurized rabbit arteries (Standen et al., 1989; Quayle et al., 1997) and capillary fragments (Langheinrich and Daut, 1997; Mederos y Schnitzler et al., 2000; Wu et al., 2006), highlighting the robust influence of K_ATP_ channels on membrane potential.

Vascular K_ATP_ channels represent ideal downstream targets for adenosine, a potent endogenous regulator of blood flow. Adenosine in the brain is primarily derived from the breakdown of ATP released from astrocytic endfeet or directly liberated from astrocytes under conditions in which there is a mismatch between tissue blood/oxygen supply and demand, such as during hypoxic/ischemic conditions (Haydon and Carmignoto, 2006; Martin et al., 2007). Since astrocytic endfeet are in intimate contact with capillaries, adenosine could conceivably have a major role in regulating K_ATP_ channels in cECs and pericytes, where *Adora2a*—the gene encoding A_2A_R—is expressed (Vanlandewijck et al., 2018). Our demonstration that adenosine stimulates K_ATP_ channels in the capillary network through the G_s_-A_2A_R-PKA signaling cascade suggests that capillaries are equipped with all the necessary molecular machinery for transducing glial-mediated stimuli into a subsequent electrical response (**Figure 4**). Consistent with this, our results revealed that K_ATP_ channels in capillary cells were similarly activated by CGRP, another endogenous vasodilator that operates through the same intracellular pathway (Nelson et al., 1990; Quayle et al., 1994). That glial-derived mediators act through capillary K_ATP_ channels to serve an important physiological function is supported by our exploration of the dilatory impact of capillary K_ATP_ channels on brain blood flow *in vivo*. These studies, in which laser Doppler flowmetry was used to measure changes in CBF induced by either exogenous pinacidil or adenosine in the absence or presence of GLIB, revealed an increase in CBF upon pinacidil or adenosine exposure that was sensitive to GLIB, suggesting that adenosine acts primarily through K_ATP_ channel activation to regulate CBF (**Figure 5**). Although difficult to determine, estimated basal adenosine levels are in the nanomolar range (Fredholm, 2010; O’Regan, 2005), and studies using intact tissue preparations have reported that, under physiological conditions, low nanomolar levels (≥5 nM) of adenosine can stimulate high-affinity A_2A_Rs (Li and Puro, 2001). Given that we used an adenosine concentration (5 μM) that is far above the estimated extracellular basal level (Fredholm, 2010; O’Regan., 2005), it is possible that the signaling pathway described here is engaged primarily under pathological conditions in which adenosine is profoundly upregulated, such as hypoxia/ischemia. However, the perivascular levels of adenosine are likely to be higher. The true physiological implications of our model must await future studies that establish an endogenous concentration of adenosine sufficient to readily increase CBF via K_ATP_ channels *in vivo*.

Surprisingly, we found modest or virtually no K_ATP_ currents in SMCs isolated from either brain pial (surface) arteries or parenchymal arterioles. This observation stands in marked contrast to a body of evidence suggesting functional expression of these channels in large cerebral arteries from different species, including rabbit, rat, and cat (Kleppisch and Nelson, 1995b; Parsons et al., 1991; Quayle et al., 1995; Zimmermann et al., 1997). Nonetheless, our findings are consistent with previous studies by us and others showing little or no effect of diverse K_ATP_ channel openers or inhibitors on pressurized murine cerebral arteries and arterioles; this contrasts with the case for the mesenteric circulation, where arterial diameter is highly impacted by these drugs (McCarron et al., 1991, Dabertrand et at., 2012; Mishra et al., 2015). These contrasting findings may reflect variability in channel expression among different species, different vascular beds, and/or location within the cerebrovascular tree. The absence of K_ATP_ channels in brain vascular SMCs supports the premise that adenosine-induced K_ATP_ channel-mediated hyperpolarization primarily originates in the capillary network and likely reflects a response to neural activity.

Collectively, our results provide robust evidence supporting the operation of a signal transduction pathway that links adenosine receptors to K_ATP_ channels in the capillary network and leads to hyperpolarization, vessel dilation, and increased CBF. This study raises the exciting possibility that K_ATP_ channels could represent a target of neural-derived adenosine that could influence CBF under physiological or pathological conditions.

## ACKNOWLEDGEMENTS

We thank T. Wellman and D. Enders for technical assistance. Research reported in this publication was supported by the National Heart, Lung, and Blood Institute of the National Institutes of Health under Award Number F32HL152576 (to N.R.K). Support was also provided by the Totman Medical Research Trust (to M.T.N.), the European Union Horizon 2020 Research and Innovation Programme (Grant Agreement 666881, SVDs@target, to M.T.N.), as well as grants from the National Institute of Neurological Disorders and Stroke (NINDS) and National Institute of Aging (NIA) (R01-NS-110656 to M.T.N.), the National Institute of General Medical Sciences (NIGMS) (P20-GM-135007 to M.T.N), and by the National Heart, Lung, and Blood Institute (NHLBI) of the NIH (R35-HL-140027 to M.T.N.).

## AUTHOR CONTRIBUTIONS

M.S. and N.R.K. designed the study; M.S., N.R.K., A.M., and T.J.H. performed research and analyzed data; M.S., N.R.K., and M.T.N. wrote the paper; D.H-E organized the presentation and edited the manuscript; and M.T.N. was the principal investigator.

## METHODS

### CONTACT FOR REAGENT AND RESOURCE SHARING

Further information and requests for resources and reagents should be directed to and will be fulfilled by the Lead Contact, Mark T. Nelson.

### EXPERIMENTAL MODEL AND SUBJECT DETAILS

All animals were used in accordance with protocols approved by the Institutional Animal Care and Use Committee of the University of Vermont. Adult (2–3 months old) male C57BL/6J and NG2-DsRed mice (Jackson Laboratories, USA) were euthanized by intraperitoneal (i.p.) injection of sodium pentobarbital (100 mg/kg), followed by rapid decapitation, or were used for *in vivo* laser Doppler flowmetry experiments. Animals were housed on a 12-h light/dark cycle with environmental enrichments and free access to food and water.

## METHODS DETAILS

### Cells Isolation

Individual cECs were obtained from C57BL/6J mouse brains by mechanical disruption of a small piece of brain somatosensory cortex (volume ~10–20 mm^3^) using a Dounce homogenizer, as previously reported (Longden et al., 2017). Briefly, tissue was homogenized in ice-cold artificial cerebrospinal fluid (aCSF) containing 124 mM NaCl, 3 mM KCl, 2 mM CaCl_2_, 2 mM MgCl_2_, 1.25 mM NaH_2_PO_4_, 26 mM NaHCO_3_, and 4 mM glucose. Debris was discarded by passing the homogenate through a 62-μm nylon mesh, and retained capillary fragments were enzymatically digested by incubating for 24 min at 37°C in an isolation solution composed of 55 mM NaCl, 80 mM Na-glutamate, 5.6 mM KCl, 2 mM MgCl_2_, 4 mM glucose and 10 mM Hepes (pH 7.3) containing 0.5 mg/mL neutral protease (Worthington), 0.5 mg/mL elastase (Worthington), and 100 μM CaCl_2_. Thereafter, the sample was incubated with 0.5 mg/ml collagenase type I (Worthington) for an additional 2 min at 37°C. The suspension was filtered, and the remaining tissue was washed to remove enzymes and triturated with a fire-polished glass Pasteur pipette. Single cells and small capillaries segments were stored in ice-cold isolation medium for use the same day (within ~5 h).

Single capillary pericytes were isolated from NG2-DsRed mouse brains by mechanical dissociation using a papain-based Neural Tissue Dissociation kit (Miltenyi Biotec) as previously described (Vanlandewijck et al., 2018). Briefly, a small piece of brain somatosensory cortex (volume ~10–20 mm^3^) was chopped into small pieces with sharp scissors, transferred to isolation solution composed of 55 mM NaCl, 80 mM Na-glutamate, 5.6 mM KCl, 2 mM MgCl_2_, 4 mM glucose and 10 mM Hepes (pH 7.3) containing enzyme 1 provide in the kit, and incubated for 17 min at 37°C. Following this step, enzyme 2 was added, mixed 10 times using a Pasteur pipette, and incubated for 12 min at 37°C. The brain suspension was then passed 10 times through a 20 G needle and incubated for an additional 10 min. Thereafter, the suspension was filtered through a 62-μm nylon mesh and stored in ice-cold isolation solution. Cells were used within ~5 h after dispersion.

Individual SMCs were obtained from pial arteries and arterioles isolated from a C57BL/6J mouse employing an enzymatic and mechanical dissociation procedure. Briefly, pial and parenchymal arteries were incubated in an isolation solution composed of 55 mM NaCl, 80 mM Na-glutamate, 5.6 mM KCl, 2 mM MgCl_2_, 4 mM glucose and 10 mM Hepes (pH 7.3) containing 0.3 mg/ml papain (Worthington) and 0.3 mg/ml dithioerythritol (Sigma-Aldrich) for 14 min at 37°C, and then transferred to a solution containing 0.67 mg/ml collagenase F, 0.33 mg/ml collagenase H (Sigma-Aldrich), and 100 μM CaCl_2_ and incubated for 5 min at 37°C. Single cells were released by triturating with a fire-polished glass Pasteur pipette and stored in ice-cold isolation medium for use the same day (within ~5 h).

### Electrophysiology

Patch-clamp electrophysiology was used to measure whole-cell currents in isolated cells. Currents were amplified using an Axopatch 200 amplifier, filtered at 1kHz, digitized at 10 kHz, and stored on a computer for offline analysis with Clampfit 10.7 software (Molecular Devices, USA). Whole-cell capacitance and series access resistance were measured using the cancellation circuity in the voltage-clamp amplifier. Electrophysiological analyses were performed in either the conventional or perforated whole-cell configuration. Patch pipettes were pulled from borosilicate microcapillary tubes (1.5-mm OD, 1.17-mm ID; Shutter Instruments, USA) and fire-polished (resistance, 3–5 MΩ). The bath (external) solution contained 80 mM NaCl, 60 mM KCl, 1 mM MgCl_2_, 0.1 mM CaCl_2_, 10 mM glucose, and 10 mM Hepes (pH 7.4). For the conventional whole-cell configuration, pipettes were backfilled with a solution containing 102 mM KCl, 38 mM KOH, 10 mM NaCl, 1 mM MgCl_2_, 1 mM CaCl_2_, 10 mM EGTA, 10 mM glucose, 10 mM Hepes and 0.1 mM ADP (pH 7.2). For perforated-patch electrophysiology, the pipette solution was composed of 10 mM NaCl, 26.6 mM KCl, 110 mM K^+^ aspartate, 1 mM MgCl_2_, 10 mM Hepes and 250–250 μg/ml amphotericin B, added freshly on the day of the experiment. Experiments were performed at negative membrane potentials (−70 mV), and the amplitude of K^+^ currents at hyperpolarized potentials was enhanced by raising external K^+^ to 60 mM. Under these conditions, K^+^ currents are inward and are detected as downward deflections. The mean capacitance of cECs, pericytes, pial and parenchymal SMCs averaged 10.40 ± 0.17 (*n* = 92), 10.36 ± 0.29 (n = 54), 15.17 ± 0.42 (n = 18) and, 11.13 ± 0.51 (n = 14) respectively. All experiments were performed at room temperature (~22°C).

### Membrane Potential Measurements

The whole-mount pressurized retina was utilized for recording from superficial capillary ECs and pericytes. Implementation of the pressurized retina for studying microvascular reactivity has been described previously (Ratelade et al. 2020, Gonzales et al. 2020). Following euthanasia, the entire orbit, including the eye, optic nerve and attached vasculature and muscles, was removed from the mouse. The retina and attached optic nerve and ophthalmic artery were dissected in 4°C Ca^2+^-free buffer containing 119 mM NaCl, 3 mM KCl, 0 mM CaCl_2_, 3 mM MgCl_2_, 5 mM glucose, 26.2 mM NaHCO_3_ and 1 mM NaH_2_PO_4_, bubbled with 95% O_2_/5% CO_2_, pH 7.4. The isolated retina (with small sclera patch intact) with attached nerve and ophthalmic artery were transferred into a perfusion chamber containing a fixed silicone platform. The ophthalmic artery was cannulated, and retina was pinned (30 μm tungsten pins) en face, using minor peripheral cuts to create a flattened surface. The entire preparation was perfused at 36°C at 5 mL/min with a bath solution containing 119 mM NaCl, 3 mM KCl, 2 mM CaCl_2_, 1 mM MgCl_2_, 5 mM glucose, 26.2 mM NaHCO_3_, 1 mM NaH_2_PO_4_ bubbled with 95% O_2_/5% CO_2_ (pH 7.4). The retina vasculature was pressurized to 50 mmHg with the same bath solution using a gravity column. After 15 min, perfusion was briefly stopped and 170 μL of papain (4 U/mL in bath solution) was added dropwise directly above the retina prep (bath volume, ~2.5 mL). Three minutes later, the papain solution was washed out by resuming perfusion. Pericytes and cECs were identified on superficial capillaries of the retina using brightfield microscopy. Cells were impaled using microelectrodes (pulled to ~200–300 MΩ and filled with a 0.5-M KCl solution containing 3 μM iFluor 488 hydrazide; AAT Bioquest). Membrane potential recordings were made using an AxoClamp-2A digital amplifier and HS-2 headstage (Molecular Devices. Signals were digitized and stored using Digidata 1322A and pClamp 9 software (Molecular Devices). Recordings satisfying the following criteria were considered successful: 1) steep negative deflection of membrane potential upon impalement; 2) steady resting membrane potential for at least 1 minute; and 3) immediate return to 0 mV upon removal of the microelectrode. Following membrane potential recordings, recorded cell type was confirmed by z-stack imaging of 488-hydrazide using an upright spinning-disk confocal microscope (also used for brightfield location of cells).

### *In vivo* Cranial Window and Laser Doppler Flowmetry

Relative CBF was continuously monitored at the site of the cranial window using laser Doppler flowmetry, as previously described (Girouard et al., 2010). Mice were anesthetized with isoflurane (5% induction, 2% maintenance). A catheter was inserted into the femoral artery for recording mean arterial pressure. A cranial window was next performed. Briefly, the skull was exposed and cleaned, and a stainless-steel head plate was attached over the left hemisphere using a mixture of dental cement and superglue. After securing the head plate to a holding frame, the somatosensory cortical territory of the brain was exposed by drilling a small circular (~2 mm diameter) section of the parietal bone of the skull. The underlying dura was carefully removed, and the cranial window was superfused with aCSF solution (37°C). After surgery, isoflurane anesthesia was discontinued and replaced with α-chloralose (50 mg/kg, i.p.) and urethane (750 mg/kg, i.p.), and the depth of anesthesia was confirmed by testing corneal reflexes and motor responses to tail pinch. CBF was continuously recorded using a laser Doppler probe (Perimed) connected to a data-acquisition system (LabChart Software; AD Instruments). The probe was positioned stereotaxically ~0.5 mm above the cortical surface at a site distant from visible pial arteries. Arterial pressure and CBF were equilibrated for 30 min before experimentation. Cerebrovascular reactivity was assessed by testing CBF responses prior to and following topical superfusion (10–30 min) of aCSF solution containing the studied pharmacological agents. Blood pressure was monitored throughout the experiment, and body temperature was measured rectally and maintained at 37°C with a feedback-controlled heating blanket.

### QUANTIFICATION AND STATISTICAL ANALYSIS

Data are expressed as means ± SEM. Where appropriate, we performed paired or unpaired t tests using Graphpad Prism 8 software to compare the effects of a given condition or treatment. P-values ≤ 0.05 were considered statistically significant, and stars in figure panels denote significant differences. Sample size estimation was predicted based on similar experiments performed previously in our laboratory. Patch-clamp and sharp electrode data were additionally analyzed using Clampfit 10.7 software (Molecular Devices).

## Supplemental Information

**Figure S1.**
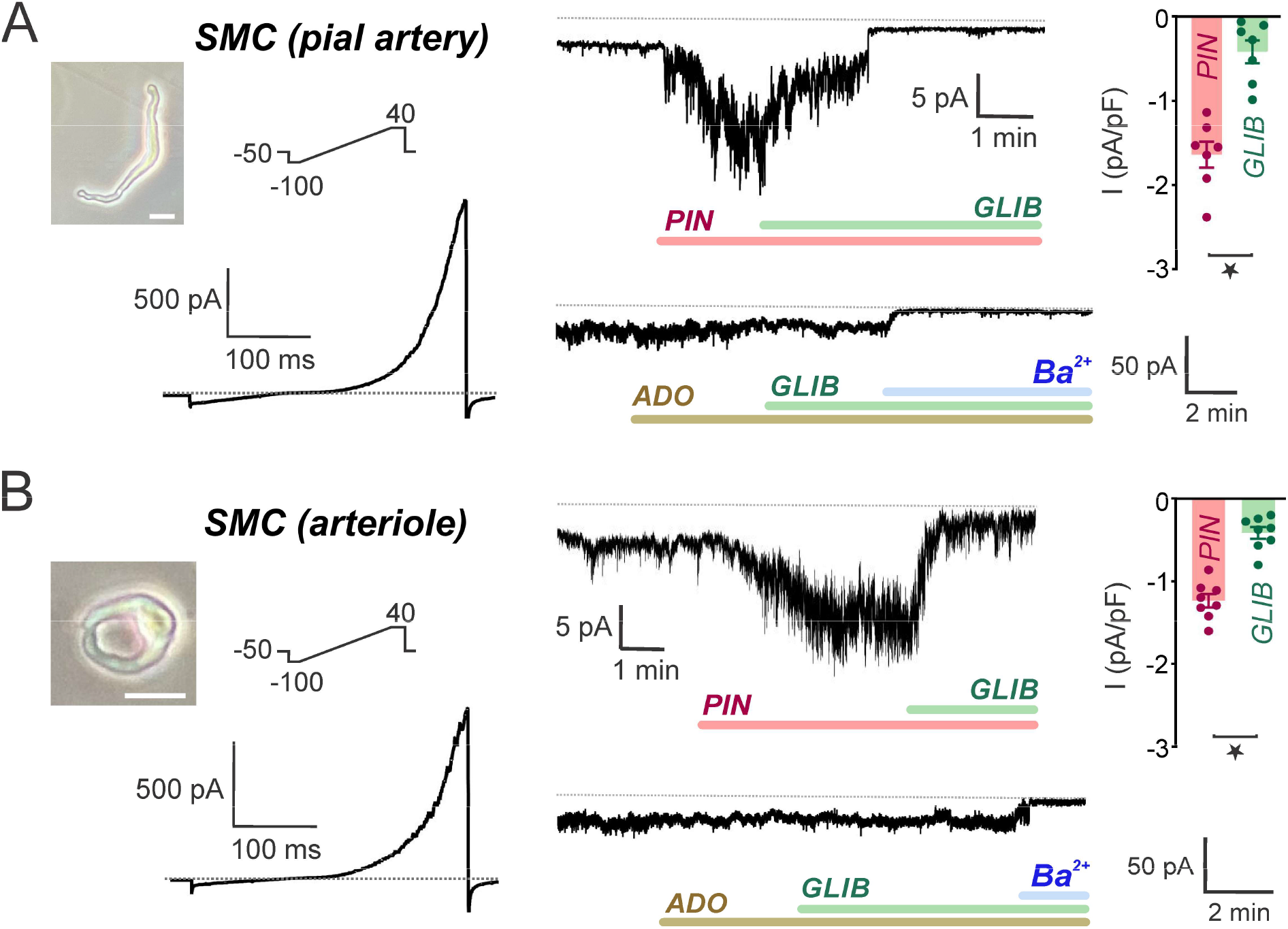
Pinacidil and adenosine evoke small (pinacidil) or no (adenosine) glibenclamide-sensitive K_ATP_ currents in SMCs from brain pial and parenchymal arteries. Representative images and current-voltage relations (*left*), original traces (*middle*) and summary data (*right*) illustrating the small effect of pinacidil (*PIN*, 10 μM; *top*) and lack of effect of adenosine (*ADO*, 25 μM; *bottom*) on whole-cell currents in freshly isolated SMCs from pial (A) (*n* = 7 and *n* = 8 for *PIN* and *ADO*, respectively) and parenchymal (B) (*n* = 8 and *n* = 6 for *PIN* and *ADO*, respectively) cerebral arteries dialyzed with 0.1 mM ATP (and 0.1 mM ADP). Summary data show *PIN*-induced current density prior to and following GLIB treatment. Further addition of BaCl_2_ (100 μM) effectively blocked inward currents at a holding potential of −70 mV. External and internal K^+^ were 60 and 140 mM, respectively. Dotted line represents 0 current level. Data are presented as means ± SEM (\**P* < 0.05, paired Student’s *t*-test). Scale bars, 20 μm.

**Figure S2.**
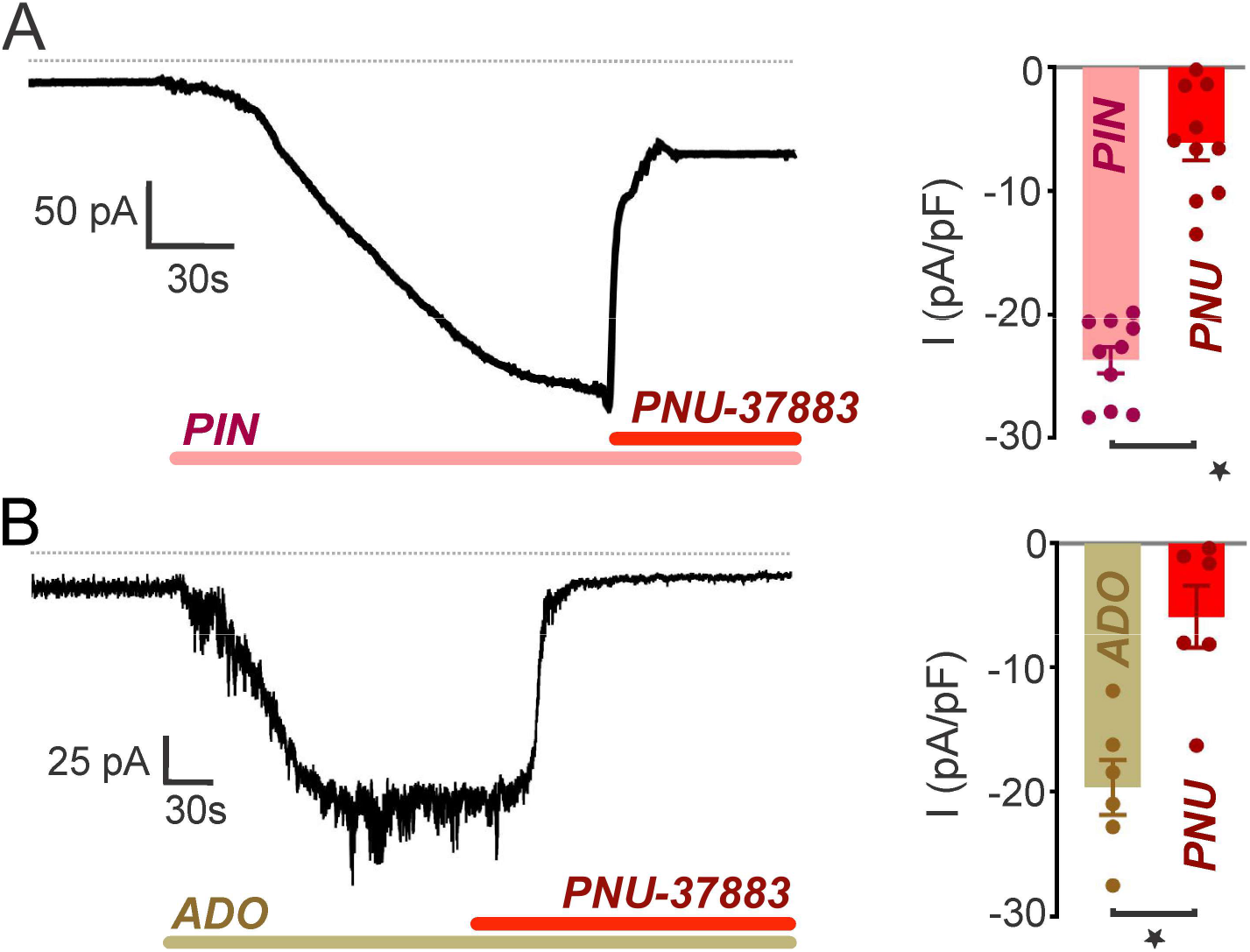
Pinacidil- and adenosine-evoked currents in cECs are sensitive to PNU-37883. Representative traces and summary data (B) showing the inhibitory effects of PNU-37883 (*PNU*; 10 μM) on currents evoked by pinacidil (A; *PIN*; 10 μM) or adenosine (B; *ADO*; 25 μM) from a holding potential of −70 mV in cECs dialyzed with 0.1 mM ATP (*n* = 10 and, *n* = 6, respectively). Summary data show pinacidil-induced current density prior to and following PNU-37883 treatment. External and internal K^+^ were 60 and 140 mM, respectively. The pipette solution also contained 0.1 mM ADP. Dotted line represents 0 current level. Data are presented as means ± SEM (\**P* < 0.05, paired Student’s *t*-test).

**Figure S3.**
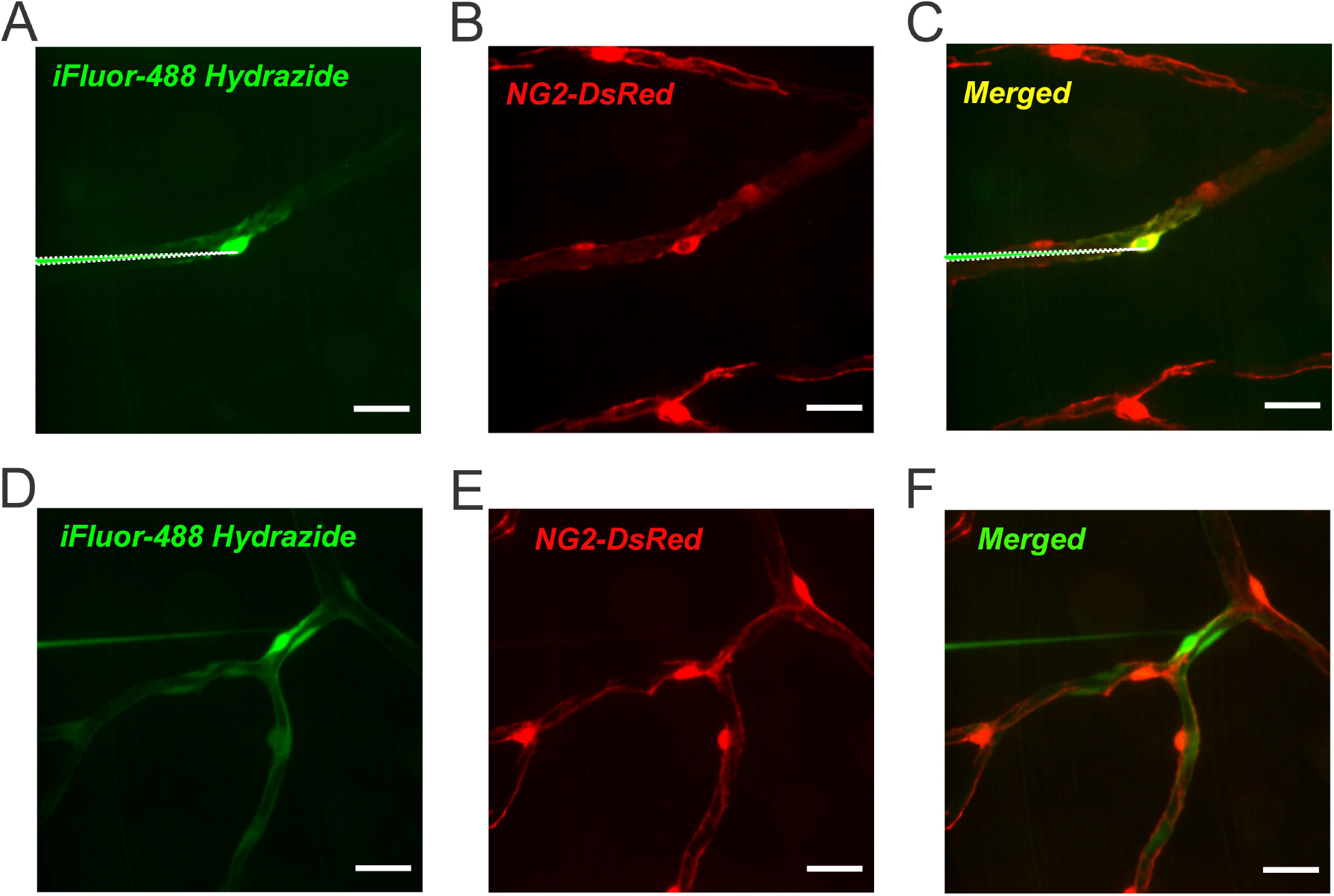
Impaled, fluorescent-hydrazide–filled pericytes and cECs were verified using *en face* pressurized retina preparations from NG2-DsRed mice. (A) Z-projection of spinning-disk confocal image showing an impaled distal pericyte filled with fluorescent hydrazide. (B) The strong NG2-DsRed fluorescence confirmed that the impaled cell was a pericyte. (C) Merged image showing color overlay of both fluorescent markers in the pericyte. Note that Z-projections were generated after removal of the microelectrode; the previously imaged position of the microelectrode was added to the images for better interpretation of impalement location. (D) Z-projection image illustrating an impaled cEC filled with fluorescent hydrazide. (E) The cell impaled was a cEC, as evidenced by its absence of NG2-DsRed fluorescence. (F) Merged image showing lack of colocalization of both fluorescent markers in the cEC. Scale bars, 20 μm.

**Figure S4.**
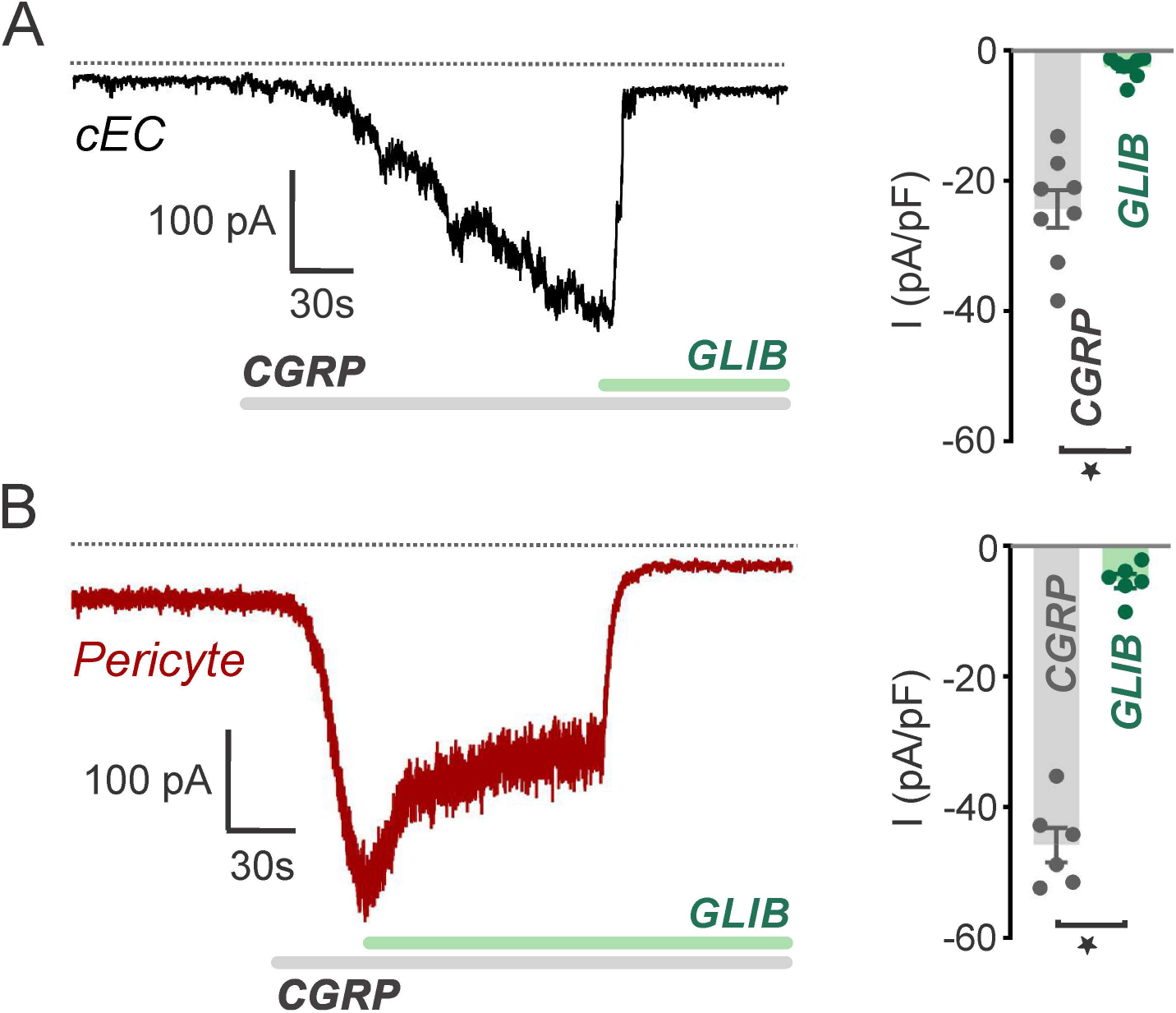
CGRP activates K_ATP_ currents in capillary EC and pericytes. Representative recordings (*left*) and summary data (*right*) illustrating glibenclamide (*GLIB*; 10 μM)-sensitive currents stimulated by CGRP (50 nM) from a holding potential of −70 mV in cECs (A) and pericytes (B) dialyzed with 0.1 mM ATP (*n* = 8 and *n* = 6 for cECs and pericytes, respectively). Summary data show CGRP-induced current density prior to and following GLIB treatment. External and internal K^+^ were 60 and 140 mM, respectively. The pipette solution also contained 0.1 mM ADP. Dotted line represents 0 current level. Data are presented as means ± SEM (\**P* < 0.05, paired Student’s *t*-test).

